# Inferring binding rates from enzymatic turnover time statistics

**DOI:** 10.1101/2025.02.04.636412

**Authors:** Divya Singh, Michael Urbakh, Shlomi Reuveni

## Abstract

We present a method to extract enzyme-substrate binding rates from observations of enzymatic turnover times. Our approach involves segregating the short-time statistics of the turnover time distribution and inferring from it the binding rate in a general and robust manner. Beyond determining binding rates, the approach developed herein also allows us to infer: (i) the fertile collision probability *p*, i.e., the probability that a collision between an enzyme and a substrate will result in the formation of an enzyme-substrate complex; and (ii) the fertile catalysis probability *ϕ*_*cat*_, i.e., the probability that the enzyme-substrate complex will lead to the formation of a product. It has long been known that *pϕ*_*cat*_ ≪ 1, indicating that most enzymes operate far from maximal efficiency, yet separating the contributions of fertile binding and catalysis was not possible in lieu of direct binding rate measurements. Our method overcomes this limitation by enabling precise inference of binding rates from turnover times, which in turn opens the door for a more detailed understanding of enzymatic efficiency.

## Introduction

Enzymatic reactions begin with the binding of a free enzyme to a substrate, forming an enzyme-substrate complex. ^1–4^ This initial step is mandatory for product formation, yet the complex can also revert back to its unbound state. Classical models of enzyme kinetics, such as the Michaelis-Menten mechanism,^5^ thus include three key kinetic transitions: binding, unbinding, and catalysis. While these transitions collectively determine the overall turnover time, their specific contributions remain obscured within the total turnover time statistics, making it difficult to isolate and analyze their individual effects. Specifically, we examine the binding rate, *k*_*on*_, a crucial yet difficult-to-measure parameter in enzyme kinetics, whose accurate determination remains a persistent challenge.^6,7^

Single-molecule fluorescence experiments have traditionally focused on product formations events, leaving the binding and unbinding steps of enzymatic catalysis largely unobserved. ^8–20^ Theoretical approaches face similar challenges, as the inherent stochasticity and convoluted nature of enzymatic reactions make it difficult to disentangle individual kinetic events from turnover time data. ^2,21–28^ Addressing these limitations often requires specialized control experiments capable of directly tracking the binding process, but such experiments are technically demanding and challenging to perform.^29–35^

Advancements in the analysis of turnover time distributions have opened new avenues for inferring hidden kinetic parameters.^13,15,17,36–39^ Specifically, it has been shown that the short-time statistics of a first-passage time, e.g., the turnover time of an enzymatic reaction, allows one to determine the number of intermediate steps in a biochemical network, probabilities associated with rare events, and also provide insight on relevant reaction timescales.^40–45^

Inspired by these developments, we ask whether short-time turnover statistics can be used to infer the binding rate, *k*_*on*_? In what follows, we will show that this is indeed possible, providing a general and robust approach to infer the binding rate from turnover time data. Prior to doing so, we will review some basic relations between the binding rate and other key observables in enzymatic catalysis. These relations will clarify that beyond characterization of the binding phase, determining the binding rate also allows one to separate the contributions of binding and catalysis to enzymatic efficiency, offering deeper insight into the overall reaction mechanism.

The binding phase of enzymatic catalysis can be described by the diffusion-limited collision rate, *k*_*d*_, and the fertile collision probability *p*, which represents the probability that a collision between an enzyme and a substrate molecule leads to binding (Fig. 1; binding phase). Consequently, the binding rate *k*_*on*_ can be expressed as

**Figure 1:**
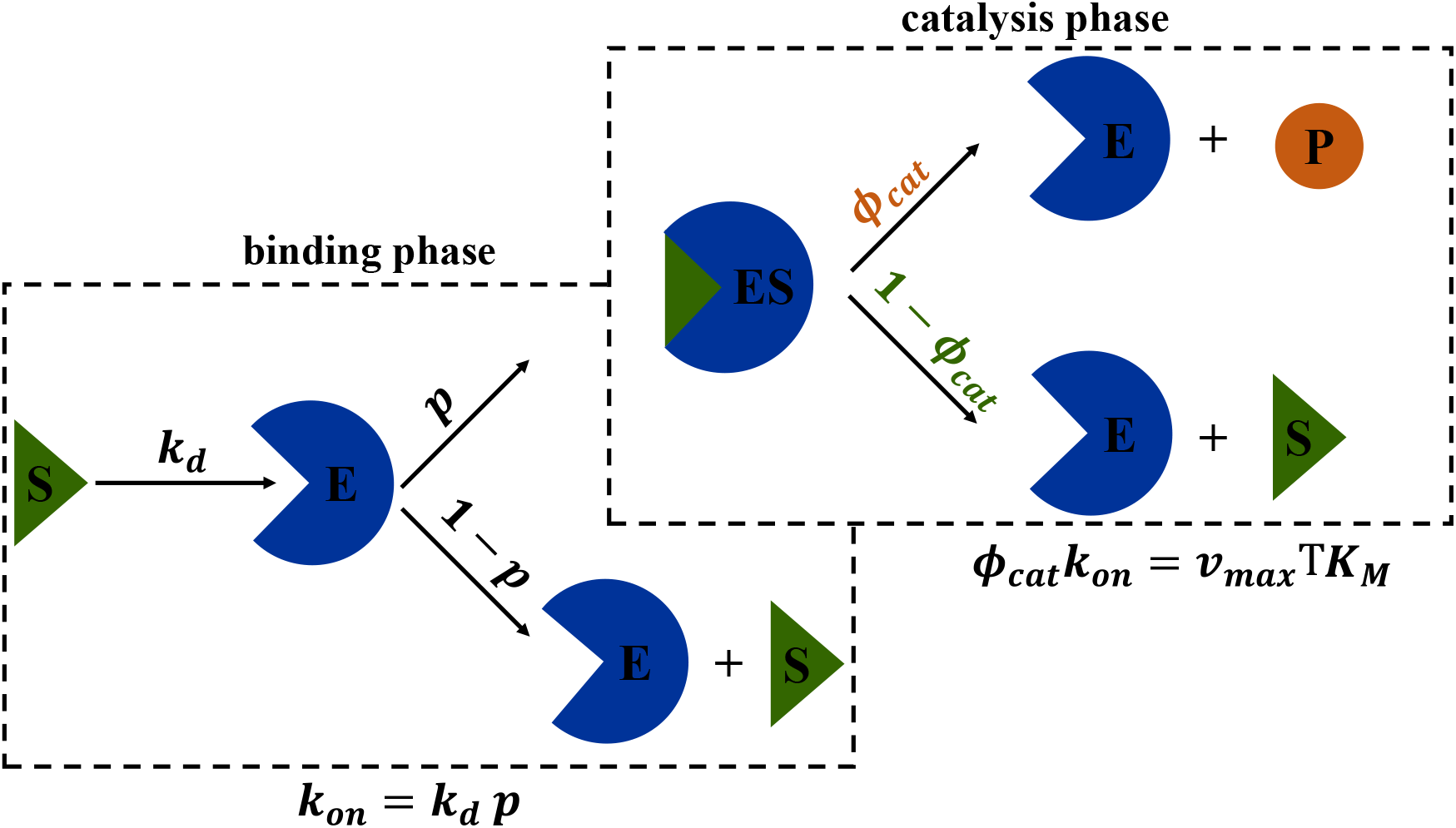
Schematic description of enzymatic catalysis. Collisions between the free enzyme *E* and a substrate molecule *S* occur at the diffusion-limited rate, *k*_*d*_. Not every collision is fertile: an enzyme-substrate complex *ES* is only formed with probability 0 *< p <* 1. Thus, the effective binding rate is *k*_*on*_ = *k*_*d*_*p*. The formation of the enzyme-substrate complex does not guarantee successful formation of a product: catalysis only occurs with probability 0 *< ϕ*_*cat*_ *<* 1. Equations (1) and (2) relate *k*_*on*_, *p*, and *ϕ*_*cat*_, which are largely unknown, to the widely available *k*_*d*_, *v*_*max*_, and *K*_*M*_ .

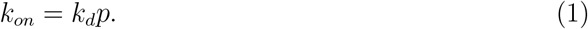

This equation encapsulates the binding process in terms of diffusion and the likelihood of productive enzyme-substrate encounters. Once bound, the enzyme may either undergo catalysis or revert to its unbound state, with probabilities *ϕ*_*cat*_ and 1−*ϕ*_*cat*_, respectively (Fig. 1; catalysis phase).

The binding rates, and fertile catalysis probabilities, of enzymes are largely unknown. Yet, these parameters can be related to two other kinetic parameters that are widely available: the (per-enzyme) maximal turnover rate, *v*_*max*_ and the Michaelis-Menten constant, *K*_*M*_ .^46,47^ The latter is equal to the substrate concentration at which half the maximal turnover rate is achieved. Indeed, it can be shown that the following relation is universal

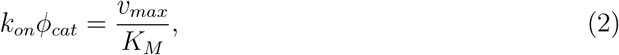

in the sense that it holds for Markovian and non-Markovian enzymes alike (SI).

Equation (2), connects the widely available catalytic efficiency (also known as the specificity constant), *v*_*max*_*/K*_*M*_ , to the underlying kinetic events, allowing one to deduce *ϕ*_*cat*_, the probability of catalysis, once *k*_*on*_ is known. Also, recalling Eq. (1), we see that given *k*_*on*_, the fertile collision probability *p* can be easily estimated since the diffusion-limited collision rate is usually known. For example, under experimental conditions where a sea of substrate molecules surrounds a tethered enzyme we have *k*_*d*_ = 4*πDR*, where *D* is the diffusion constant of the substrate molecules and *R* is roughly the radius of the enzyme.^48,49^

Most enzymes operate well below the diffusion limit, with product formation occurring in fewer than one in 10^4^ reaction trajectories due to inefficiencies in both binding and catalysis.^46,47^ These inefficiencies are captured by *pϕ*_*cat*_ ≪ 1. Yet, separating the contributions of binding and catalysis to this product has so far been challenging. Our method addresses this gap, enabling inference of the enzyme-substrate binding rate, which in turn allows for the disentanglement of *p* and *ϕ*_*cat*_.

In what follows, we will present a theoretical framework for extracting binding rates from turnover time data. Our starting point is the turnover-time distribution of the generalized Michaelis-Menten model, which has been derived in Singh et al.^50^ By focusing on short-time statistics, we isolate the binding rate *k*_*on*_ and demonstrate how it can be inferred from data. We validate our approach, through numerical simulations of various enzymatic systems, showing that inferred binding rates closely match their theoretical values. Lastly, we discuss the broader implications of these findings for enzymatic kinetics and potential applications.

## Inferring binding rates from enzymatic turnover time statistics

In a recent study, we derived a detailed functional form of the turnover-time distribution for a generalized Michaelis-Menten reaction mechanism.^50^ To describe this result, we introduce some basic notation.

First, observe that any enzymatic turnover cycle begins with the binding of a substrate molecule to a free enzyme. Thus, following a stochastic binding time, *T*_*on*_, an enzyme-substrate complex, *ES*, is formed. The *ES* complex may proceed to catalysis, leading to the formation of a product molecule and the regeneration of the free enzyme (*E* + *P* ), or it may revert to the unbound state *E* + *S* without forming a product.

Next, we note that the conditional distribution of the waiting time in the *ES* state given that unbinding occurs, need not be the same as the conditional distribution of the waiting time given that catalysis occurs. In other words, enzyme dynamics can have non-Markovian characteristics. Letting *ϕ*_*cat*_ stand for the probability that catalysis occurs before unbinding, we denote the conditional waiting time in this scenario by 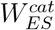, and the probability density function that describes this random variable by 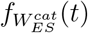. The probability that unbinding occurs before catalysis is then 1 − *ϕ*_*cat*_, and we denote the conditional waiting time in this scenario by 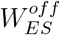, and the probability density function that describes this random variable by 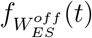. In Fig. 2(a), we illustrate these random times graphically considering several turnover trajectories for example.

**Figure 2:**
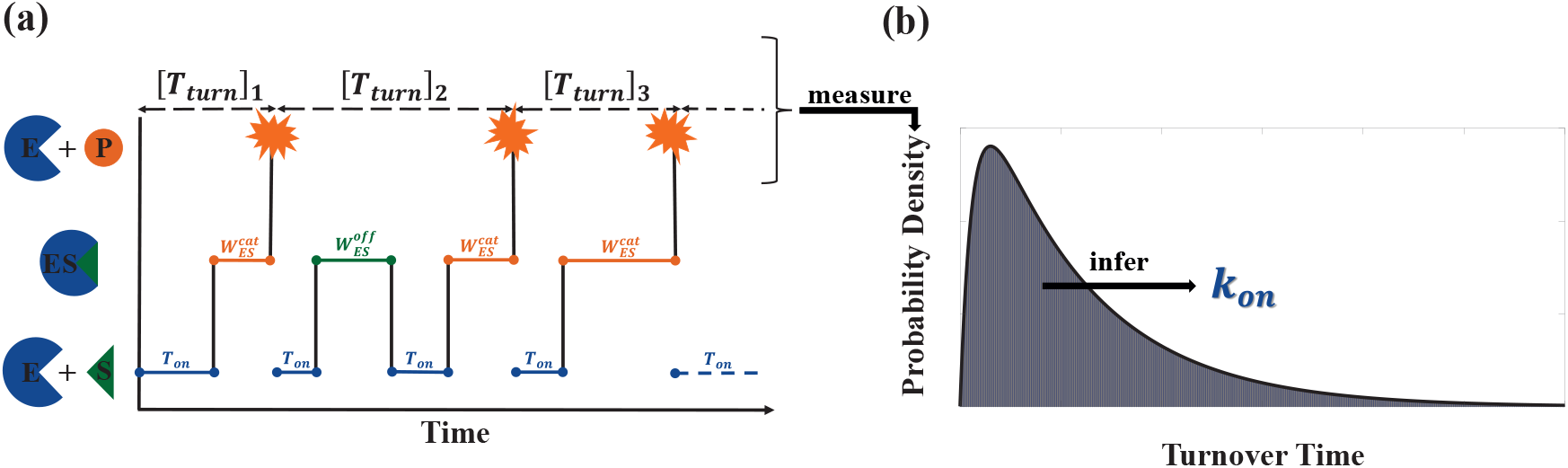
Inferring binding rates from enzymatic turnover time statistics. (a) A schematic representation of turnover trajectories, showing three consecutive enzymatic product formation events. The first turnover time, [*T*_*turn*_]_1_, is a sum of the substrate binding time, *T*_*on*_, and the conditional waiting time for catalysis, 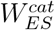. After the first turnover event, the enzyme *E* returns to its free state, binds a new substrate *S*, and forms the enzyme-substrate complex *ES*. Following a conditional waiting time for unbinding, 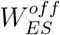, this complex breaks apart. An additional binding event, followed by a catalysis event, are then required to form a product. The times of these events tally to give the second turnover time [*T*_*turn*_]_2_. The third turnover time, [*T*_*turn*_]_3_, includes a single binding and catalysis events. Note that the binding time, and the conditional waiting times for unbinding and catalysis, are all random. (b) The turnover time distribution is reconstructed from multiple turnover events. The short-time behavior of this distribution can then be used to infer the binding rate, *k*_*on*_.

A complete distribution of the turnover time can be constructed by observing multiple turnover events (Fig. 2(b)). Embedded within this distribution is valuable information about the enzyme-substrate binding rate, *k*_*on*_, which governs the enzyme-substrate association rate. However, this information is obscured by the stochastic nature of the enzymatic cycle as we discuss next.

We model enzymatic catalysis by assuming that substrate binding times are exponentially distributed with rate *k*_*on*_[*S*], where [*S*] is the substrate concentration. In contrast, the waiting times associated with catalysis and unbinding are represented by arbitrary probability distributions. This level of generality allows us to capture the underlying kinetics without needing to specify microscopic details, which makes this approach broadly applicable to a wide range of enzymatic systems.

In Singh et al., ^50^ we showed that the Laplace transform of the turnover-time distribution 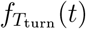, can be written in terms of the times and probabilities defined above

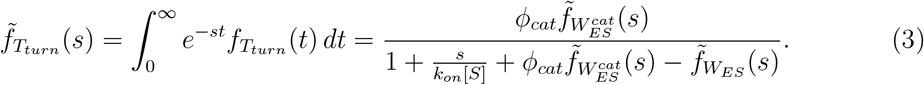

Here, 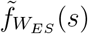 is the Laplace transform of the unconditional waiting time distribution in the *ES* state. This quantity is (by definition) a weighted sum of the Laplace transformed conditional waiting time distributions 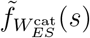 and 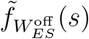, with weights *ϕ*_cat_ and 1 − *ϕ*_cat_, respectively. It can thus be written as: 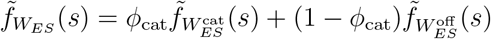.

Equation (3) provides a direct link between observable turnover time data and the enzymatic binding rate, *k*_on_. Yet, inference of the binding rate is not straightforward. The primary challenge lies in the fact that the probability density function (PDF) of turnover times incorporates contributions from a wide range of trajectories, some of which involve multiple binding and unbinding events (see Fig. 2(a)). These repeated events increase the complexity of the turnover-time statistics.

To simplify the analysis, it is advantageous to isolate, or at least amplify, the contribution of trajectories involving only a single binding and catalysis step, and no unbinding events (e.g., trajectories 1 and 3 in Fig. 2(a)). Such trajectories dominate the short-time behavior of the turnover-time PDF, as the inclusion of unbinding events typically increases the turnover time. To achieve this, we introduce an exponentially distributed cutoff time, *T*^*^, whose probability density is given by

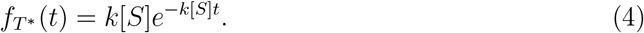

Here, *k*[*S*] is a ‘censoring’ rate that is proportional to the substrate concentration. For clarity, the substrate concentration [*S*] is given by the enzymatic system at hand, while *k* is a control variable that regulates the censoring rate, *k*[*S*]. By tuning *k*, one can ensure that cutoff times are predominantly short, allowing focus on the corresponding turnover events.

Next, we define a conditional random variable, 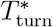, which represents turnover times conditioned on being smaller than *T*^*^. We thus have

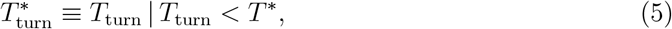

as the turnover time censored by the exponentially distributed cutoff time *T*^*^. In the Supplementary Information (SI), we derive the distribution of 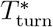, and show that its Laplace transform is given by

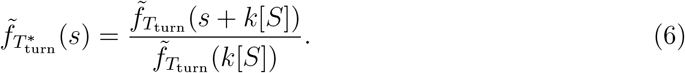

From here, we compute the first moment of 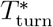, which is given by

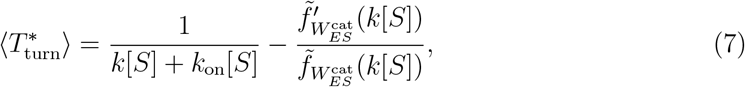

where 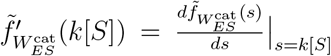. We will now show that this formulation enables us to extract the binding rate *k*_on_ by leveraging the short-time behavior of the turnover-time distribution. The use of *T*^*^ as a cutoff effectively amplifies the contribution of simple binding-and-catalysis trajectories, ensuring that the inferred *k*_on_ is robust and less influenced by complex, multi-step pathways.

To proceed, we note that the second term on the right-hand side of Eq. (7) can be simplified for a wide class of probability density functions (PDFs) that exhibit a power-law behavior, 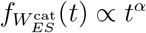, at short time scales. Notably, the Exponential, Gamma, and Weibull distributions all belong to this class. In the Supplementary Information (SI), we show that for distributions of this form, and when *k*[*S*] ≫ 1, Eq. (7) reduces to

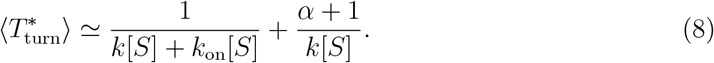

Equation (8) enables the determination of *k*_on_ in two steps. First, *α* is extracted by considering 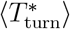 at large values of the censoring parameter (*k* ≫ *k*_on_), where we have

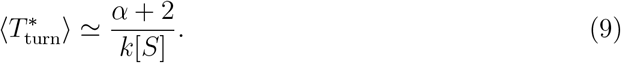

The value of *α* can then be determined by fitting Eq. (9) to data. Once the power-law exponent *α* is known, the asymptotic expression in Eq. (8) can be rearranged to yield

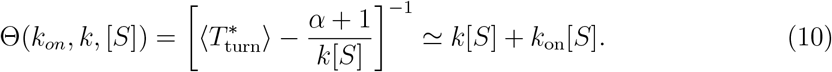

Plotting the left-hand side of Eq. (10) against the censoring rate, *k*[*S*], allows one to determine the binding rate *k*_on_ by fitting a straight line. The intercept, divided by the substrate concentration [*S*], gives the desired binding rate.

## Extracting the binding rate from data: Markovian Case Study

In Fig. 3, we demonstrate how to infer the substrate binding rate *k*_*on*_ from turnover time data. To this end, we consider a Markovian enzyme as a representative example. Specifically, we assume that binding, unbinding, and catalysis are all exponential processes with rates *k*_*on*_ = 5 *µM*^−1^*s*^−1^, *k*_*off*_ = 10 *s*^−1^ and *k*_*cat*_ = 20 *s*^−1^. These parameters give a Michaelis-Menten constant of 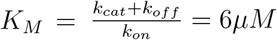, and we set [*S*] = 5*K*_*M*_ .

**Figure 3:**
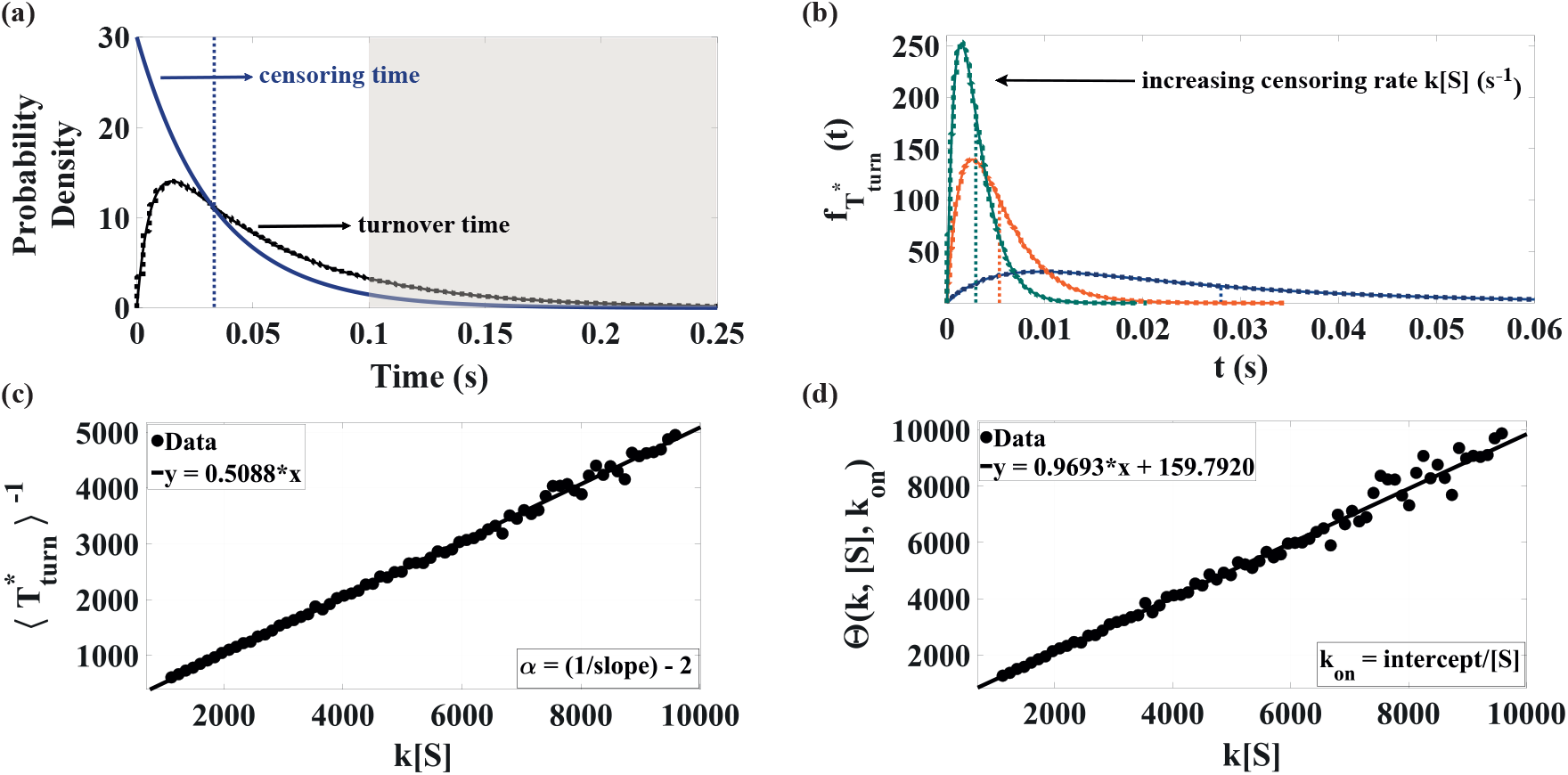
Inferring the binding rate step by step. (a) Turnover-time probability distribution for a Markovian enzyme (black squares: simulation data; solid black curve: theory; parameters given in text), and the distribution of the exponential censoring time from Eq. (4) (solid blue line, parameter given in text). The mean censoring time ⟨*T*^*^⟩ is shown as a vertical dashed blue line. Turnover times significantly exceeding this value (gray shaded area) are effectively censored. (b) Distribution of the conditional (censored) turnover-time from Eq. (5) for [*S*] = 30 *µM* and three values of the control parameter *k* = 1, 20, 30 *µM*^−1^*s*^−1^ (blue, orange and green symbols, respectively). The symbols, which come from numerical censoring of the turnover data in panel (a), are corroborated by theoretical results (solid lines, see SI). Vertical dashed lines indicate the mean value of each distribution. (c) Estimating the power-law exponent *α* by fitting a straight line to the data points gives *α* ≃ −0.0346, which is close to the theoretical value of zero. (d) Inference of *k*_*on*_. The left-hand side of Eq. (10) is plotted against *k*[*S*] and a linear fit yields an estimate for the binding rate. We find *k*_*on*_ ≃ 5.32, which closely matches the true value.

Each panel in Fig. 3 corresponds to a specific step in the inference procedure. We start with Fig. 3(a), which depicts the turnover-time probability distribution for forming a single product molecule. Here, substituting for experimental data, an estimate for this distribution is obtained by simulating the Markovian model defined above, generating 10^7^ turnover events (symbols). This estimate is corroborated using the analytical turnover time distribution which is known for the Markovian case (solid black line). To extract the binding rate from the simulated data, we introduce an exponential censoring-time whose distribution, *f*_*T*\*_ (*t*), is given by Eq. (4), with [*S*] = 30 *µM* , and *k* = 1 *µM*^−1^*s*^−1^ that is set here for illustration purposes. The mean censoring time, which is given by ⟨*T*^*^⟩ = (*k*[*S*])^−1^ = 1*/*30 *s*, is indicated by the dashed blue line. Turnover times that significantly exceed this cutoff are effectively discarded (gray shaded area), allowing us to focus on short turnover time trajectories.

In Fig. 3(b), we plot distributions of the censored turnover time, 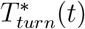, that was defined in Eq. (5). These distributions are plotted for three values of the control parameter, *k* = 1, 10, 20 *µM*^−1^*s*^−1^, and they are obtained from the empirical turnover time distribution in Fig. 3(a) via a simple numerical sampling procedure (see SI). Dashed vertical lines mark the means of the corresponding distributions. Results obtained numerically are corroborated using the analytical distributions of the censored turnover times (solid lines, see SI).

Next, we repeat the procedure illustrated in Fig. 3(b), calculating the average censored turnover time, 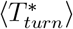, for increasing values of the censoring parameter *k*. In Fig. 3(c), we plot 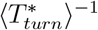 against *k*[*S*]. For *k*[*S*] ≫ 1 and *k* ≫ *k*_*on*_, we have from Eq. (9): 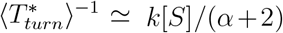. This allows us to extract *α* by fitting the corresponding region of the data with a straight line. Doing so we find *α* ≃ −0.0346, which is very close to the theoretical value of *α* = 0 that is expected for the exponential catalysis time distribution in the Markovian Michaelis-Menten scheme (see SI).

Finally, Fig. 3(d) illustrates the inference of *k*_*on*_ using Eq. (10). The left-hand side of Eq. (10), 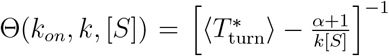, is plotted against *k*[*S*], and a linear fit to the data is made. The intercept of the fitted line, divided by the substrate concentration [*S*], yields *k*_*on*_ ≃ 5.32 ± 1.31. This estimate is consistent with theoretical value *k*_*on*_ = 5.

## Summary and Discussion

Single-molecule measurements provide powerful tools for investigating the microscopic mechanisms underlying enzymatic reactions. Yet, these measurements have primarily focused on tracking product formation, making it difficult to disentangle the contributions of individual kinetic events within the enzyme’s molecular machinery. In this study, we introduced a method for extracting enzyme-substrate binding rates from such turnover time observations. By isolating the short-time statistics of the turnover time distribution, we directly infer the binding rate, *k*_on_, overcoming the limitations of previous approaches.

Beyond determining the binding rate, our method addresses a long-standing challenge in enzymatic kinetics by enabling the separate resolution of the fertile collision probability, *p*, and the fertile catalysis probability, *ϕ*_cat_. These probabilities quantify the likelihood of productive enzyme-substrate encounters and successful product formation, respectively. Traditionally, only their product, *pϕ*_cat_, was accessible, leaving their individual contributions indistinguishable without binding rate information. By inferring the binding rate from turnover time data, our approach overcomes this limitation, offering a more detailed picture of enzyme catalysis.

The progress made in this work complements results presented in our recent publication.^50^ There, we derived a set of high-order Michaelis-Menten equations revealing universal linear dependencies between unique combinations of turnover time moments and the reciprocal of the substrate concentration. While these equations established relationships between key kinetic parameters, they were incomplete: requiring external and independent knowledge of at least one of the unknown parameters to infer the others. The method developed here resolves this limitation by providing orthogonal means to determine the binding rate. Combined with the framework presented in Singh et al., ^50^ it provides a complete toolkit for inferring all kinetic parameters governing enzymatic reactions. This paves the way for a deeper and more nuanced understanding of enzymatic efficiency and the physical constraints underlying enzymatic catalysis.

## Acknowledgement

This project has received funding from the European Research Council (ERC) under the European Union’s Horizon 2020 research and innovation program (Grant agreement No. 947731).

